# Movement quality moderates the effect of spatially congruent cues on the stability of symmetric and asymmetric rhythmic bimanual finger movements

**DOI:** 10.1101/2024.03.18.585605

**Authors:** Ronan Denyer, Lara A. Boyd

## Abstract

Spatially congruent cues increase the speed of bimanual reach decisions compared to abstract symbolic cues, particularly for asymmetric reaches. Asymmetric rhythmic bimanual movements are less stable than symmetric rhythmic movements, but it is not well understood if spatially congruent cues similarly increase the stability of asymmetric rhythmic bimanual movements. To address this question, in *Experiment 1*, participants performed symmetric and asymmetric bimanual rhythmic finger tapping movements at different movement frequencies in time with flickering spatially congruent and abstract symbolic stimuli. As expected, symmetric movements were more stable. Spatially congruent cues similarly increased the stability of symmetric and asymmetric movements compared to abstract symbolic cues. The benefits of spatial congruence and movement symmetry were restricted to high movement frequencies (>2 hertz). To better understand if the emergence of these effects at high movement frequencies was driven by a change in movement strategy, in *Experiment 2*, video of the hands was concurrently recorded during task performance. Markerless motion tracking software revealed that participants switched from discontinuous to continuous movement strategies with increasing movement frequency. Since discontinuous and continuous movements are thought to be controlled by distinct neuro-cogntive systems, this might explain why the beneficial effects of spatial congruence and response symmetry emerged only at high movement frequencies. Overall, results from the current study indicate that the perceptual quality of the stimulus use to cue movement frequency can have powerful effects on the stability of rhythmic bimanual movements, but that these effects may depend on whether discontinuous or continuous movement strategies are selected.

## Introduction

Deriving principles of bimanual coordination has most frequently been pursued through the study of rhythmic bimanual movements. A cardinal finding from this paradigm is that mirror-symmetric bimanual movements are executed with greater stability than asymmetric movements (for reviews, see (Donchin & de Oliveira, 2004; Kelso, 1995; Swinnen, 2002; Swinnen & Wenderoth, 2004)). Less stability in asymmetric movements may be due to “crosstalk” between the neural processes responsible for translating task goals and external stimuli into rhythmic action (Cattaert et al., 1999; Marteniuk et al., 1984; Sherwood, 1994; Spijkers & Heuer, 1995; Swinnen et al., 1991). According to this view, stabilization of asymmetric bimanual movement patterns requires that sources of interference are successfully suppressed.

There are competing perspectives regarding the dominant source of neural crosstalk driving the instability of asymmetric rhythmic bimanual movements. The “motor outflow hypothesis” argues that interference within motor programming processes are the dominant source of neural crosstalk driving asymmetric instability (Donchin & de Oliveira, 2004; Serrien et al., 2002, 2003). This view is supported by transcranial magnetic stimulation (TMS) studies, which show that during ongoing performance of unimanual and bimanual rhythmic movements, corticospinal excitability is modulated in a phasic manner correlating with the rhythmic phase of the moving limb(s) (Carson et al., 2004). According to the motor outflow hypothesis, limitations in the control of asymmetrical movements can be attributed to “hard-wired” connections between motor areas of the left and right hemispheres. Mediated by the corpus callosum, asymmetric patterns of excitability of effector representations within bilateral primary motor cortices (M1) destabilize asymmetric bimanual behaviour (Carson, 2005; Eliassen et al., 2000; Franz et al., 1996).

In contrast, according to the “central processing hypothesis”, cognitive-perceptual processes upstream of motor programming, are the dominant source of neural interference driving the instability of asymmetric bimanual movements. This view is supported by evidence from discrete bimanual reaching paradigms, in which participants were required to quickly perform asymmetric or symmetric reaches after the onset of a visual cue. The target bimanual reaches were cued either symbolically by letters or cued in a spatially congruent manner through illumination of the endpoint locations. Bimanual reaches were slower in the symbolic cue condition. Importantly, symbolic cues were also associated with an increase in reaction time (RT) for asymmetric compared to symmetric reaches, while there was no difference in RT between symmetric and asymmetric reaches when spatially congruent cues were used (Diedrichsen et al., 2001, 2003, 2006; Hazeltine et al., 2003). These results suggest that constraints on planning discrete asymmetric bimanual movements are primarily driven by interference within central cognitive processes responsible for translating task-relevant information from sensory representations to action plans. Complementary neuroimaging evidence suggests that spatially congruent cue conditions facilitate use of a rapid automatic response selection system, enabling the planning of two reach trajectories in parallel. In contrast, abstract symbols force the use of a slower and serial response selection process, which introduces a potent source of interference (Diedrichsen et al., 2006).

Cognitive-perceptual factors have been shown to affect rhythmic bimanual control. For example, asymmetric index finger flexion-extension oscillations are performed with similar accuracy to symmetric oscillations if the wrist position of one hand is rotated such that the perceptual consequences of asymmetric movements are made symmetrical (Mechsner et al., 2001). Furthermore, asymmetric bimanual rhythmic finger circling is more stable when cued with flashing light emitting diodes (LEDs) that are spatially congruent with the direction of motion (Byblow et al., 1995, 1999), while asymmetric rhythmic hand circling is more easily learned and retained when participants are provided with enhanced online visual feedback of the position of their hands (Swinnen et al., 1997). Similarly, switching from symmetric to asymmetric rhythmic circling patterns results in fewer errors when the timing of switches are cued with spatially congruent stimuli cues (Wenderoth et al., 2009).

Existing investigations into the effect of spatially congruent cues on rhythmic bimanual movements primarily focus on rhythmic circling movements, whereas repetitive finger tapping movements have received less attention. This is an important distinction because rhythmic bimanual tapping and circling may be fundamentally different classes of movement with distinct coordination principles (R. Ivry et al., 2004; R. B. Ivry et al., 2002). For example, performance on rhythmic finger tapping and rhythmic circle drawing tasks does not correlate within subjects (Pope & Studenka, 2019; Robertson et al., 1999; Zelaznik et al., 2000, 2002, 2005). Furthermore, callosotomy patients cannot spatiotemporally couple hands during rhythmic bimanual circle drawing (Kennerley et al., 2002) but can spatiotemporally couple fingers when tapping in rhythm at regular frequencies (Franz et al., 1996; R. B. Ivry & Hazeltine, 1999; Tuller & Kelso, 1989). However, this latter capacity is abolished when patients are instructed to perform smooth continuous finger oscillations instead of a series of discrete taps separated by a pause (R. Ivry et al., 2004; Kennerley et al., 2002). Interestingly, patients with cerebellum damage show the opposite pattern of deficits (R. B. Ivry et al., 2002; Spencer et al., 2003). Taken together, these results suggest that rhythmic bimanual movements that are discrete rely on event timing mechanisms mediated by the cerebellum. In contrast, rhythmic bimanual movements that are continuous rely on “emergent timing” mechanisms mediated by transcallosal cortical networks with spatiotemporal coupling emerging via the control of higher order kinematic parameters such as speed (R. Ivry et al., 2004; R. B. Ivry et al., 2002; Kennerley et al., 2002).

In the current study, across two experiments we tested (1) whether cueing discrete bimanual key presses with spatially congruent visual stimuli versus abstract symbolic visual stimuli decreases choice RT (*Experiment 1*), (2) whether cueing the movement frequency of asymmetric bimanual rhythmic finger tapping movements with spatially congruent visual stimuli versus abstract symbolic visual stimuli influences the stability of bimanual spatiotemporal coupling (*Experiment 1*), and (3) whether the movement strategy used to perform rhythmic bimanual key presses changes with increasing movement frequency (*Experiment 2*). In *Experiment 1*, we assessed whether the benefits of spatially congruent cues versus abstract symbolic cues are shown when performing discrete bimanual finger movements. Participants completed two versions of a bimanual 4-choice RT task, which differed in terms of whether choices were triggered by spatially congruent or abstract symbolic visual stimuli (**Figure 1**). Reaction times for symmetric and asymmetric bimanual finger movements were recorded. Based on findings from bimanual reaching paradigms (Diedrichsen et al., 2001, 2006), we predicted that RT would be faster when movement was cued with spatially congruent stimuli (Hypothesis 1A), that RT would be faster when symmetric responses were required (Hypothesis 1B), and that the cost to RT for asymmetric responses would be significantly reduced when movement was cued with spatially congruent stimuli (Hypothesis 1C).

**Figure 1.**
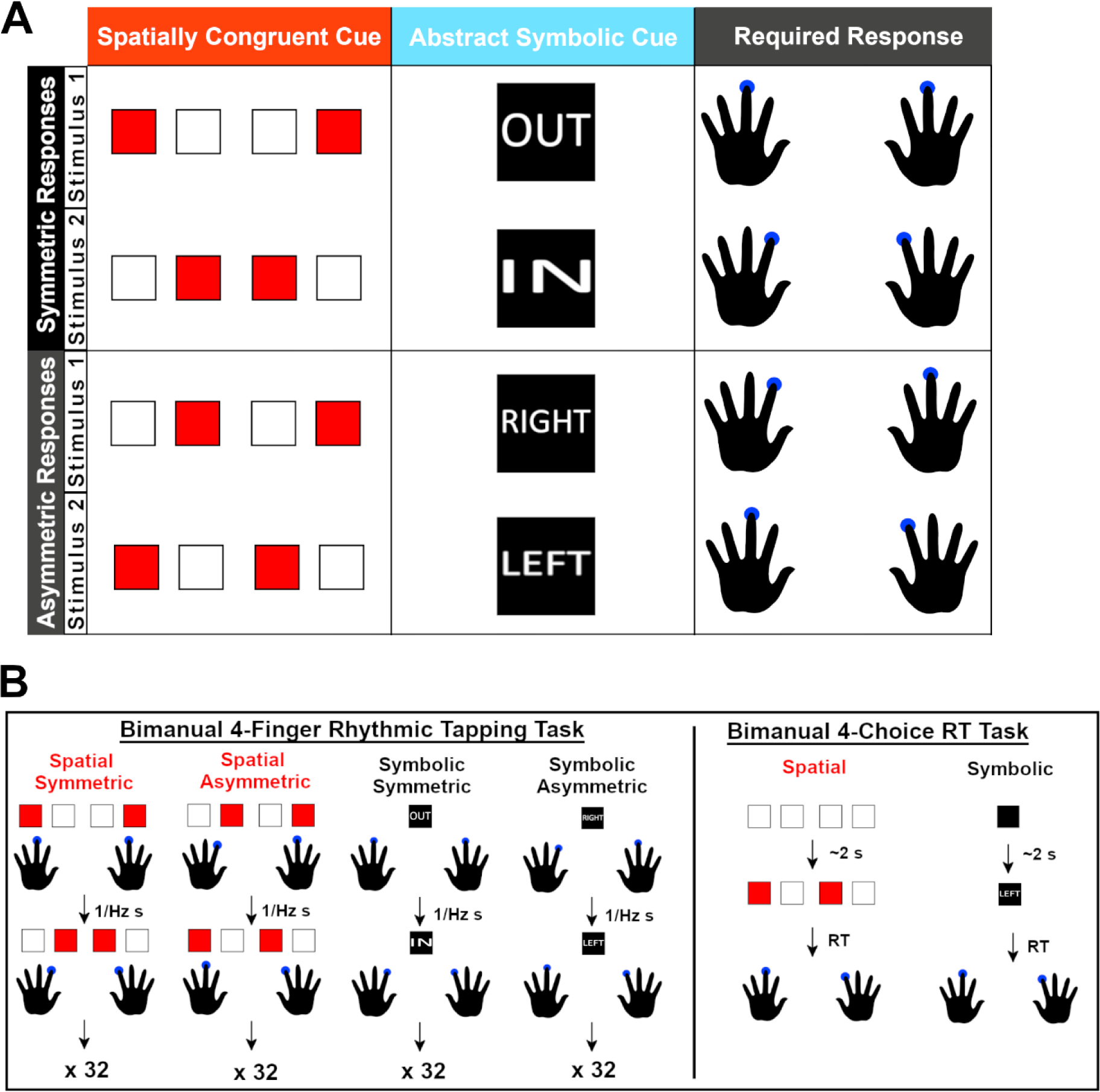
Stimulus-response mappings and tasks outline. **(A)** In both tasks, movement timing was either cued with spatially congruent visual stimuli, where red boxes directly denoted the relative spatial location of the fingers required to move, or abstract symbolic stimuli where words abstractly denoted the identity of the fingers required to move. Required responses were either symmetric or asymmetric. In the bimanual 4-finger rhythmic tapping task, participants performed 4 versions of the task, in which movement frequency was cued with either spatially congruent or abstract symbolic visual cues, and the required response pattern was either symmetric or asymmetric. **(B; left)** Participants performed 12 blocks of 32 bimanual taps for each task condition. Movement frequency started at 1 hertz (Hz) and increased by 0.2 Hz after each block. **(B; right)** Participants performed 2 versions of the bimanual 4-choice RT task, one in which the visual cues used to cue movement were spatially congruent and one in which the visual cues used to cue movement were symbols. Participants performed 60 RT trials each for the spatial and symbolic versions of the task.

In *Experiment 1*, we also assessed whether spatially congruent and abstract symbolic cues have an effect on the accuracy and stability of rhythmic bimanual finger tapping movements. Participants performed four versions of a 4-finger bimanual rhythmic tapping task. The task versions differed in terms of whether repeated symmetric or asymmetric responses were required and whether movement frequency was cued by repeating spatially congruent or symbolic visual stimuli (**Figure 1**). Rhythmic tapping was assessed across a range of movement frequencies. It was predicted that the accuracy and stability of rhythmic tapping across all task versions would decrease with increasing movement frequency (Hypothesis 2A). It was further predicted that symmetric movement patterns would be more stable compared to asymmetric movement patterns (Hypothesis 2B) and that movement patterns cued with spatially congruent cues would be more stable compared to those cued with abstract symbolic cues (Hypothesis 2C). We also hypothesized that the cost of asymmetric patterns on stability would be greater when movement patterns were cued with abstract symbolic stimuli compared to spatially congruent stimuli (Hypothesis 2D).

Whether rhythmic bimanual movements are discrete or continuous can dramatically alter the coordination principles at play. Since measures of task performance were derived entirely from key press timing data in *Experiment 1*, it was not possible to interrogate potential shifts in movement strategy from repeated discrete movements to continuous movement with data from *Experiment 1.* Therefore, to better understand how changes in movement strategy across movement frequencies might influence results from *Experiment 1,* a follow up *Experiment 2* was conducted. Participants performed the same 4-finger bimanual rhythmic tapping tasks from *Experiment 1* in a laboratory setting, but with concurrent video recording of their fingers during task performance. Recordings of finger movements were analyzed with deep learning algorithms (Mathis et al., 2018) to capture 2-dimensional spatiotemporal kinematics of the fingers during task performance. The mean relative speed and acceleration of finger trajectories in a prescribed time window around the time of key presses was used as a proxy measure of movement smoothness. It was predicted that mean relative speed would increase while mean relative acceleration would decrease as movement frequency was increased, in line with a change in movement quality from rapid discrete movements separated by long pauses to smoother continuous movement with no pauses (Hypothesis 4A). It was further predicted that changes in mean relative speed and acceleration would be similar across task conditions (Hypothesis 4B).

## Materials and methods

### Experiment 1

#### Participants

Individuals were automatically excluded if they were not aged 18-65 yr, were not fluent English speakers, or had a history of brain injury/disease. A total of 250 participants completed Experiment 1. Collection of this large sample size was facilitated by use of online platforms, which was necessary to overcome restrictions on research imposed by COVID-19. The final sample size was multiple times larger than any previous investigation using similar behavioural methods, providing more than adequate statistical power to test the described hypotheses. The majority of participants were recruited via a University of British Columbia undergraduate human research participants pool and received academic credit for participating. The remaining participants were recruited through personal relationships. All participants had the option to enter their name into a lottery for a retail store gift voucher award. Of the 250 participants, 9 did not perform key presses as required on certain task blocks and their data were excluded from the final analyses, leaving a total of 241 participants (22.1 ± 6.2 years old; 186 female, 55 male; 225 right-handed, 16 left-handed). All protocols were approved by Behavioural Research Ethics’ Board at the University of British Columbia. Before participating, individuals provided written informed consent in accordance with the Declaration of Helsinki.

#### Experimental design

Using a within-subjects design, all participants first performed two versions of a 4-finger bimanual choice RT task: one in which responses were cued with abstract symbolic stimuli (i.e., words which described the identity of the requisite bimanual finger pair), and another in which responses were cued with spatial stimuli (i.e., boxes which denoted the relative spatial location of the requisite bimanual finger pair; **Figure 1**). Next, participants performed four versions of a 4-finger bimanual rhythmic tapping task. Task versions differed in terms of whether symmetric or asymmetric tapping patterns were required and whether movement timing was cued with abstract symbolic or spatially congruent stimuli (**Figure 1)**. The order of task stimulus type (symbolic/spatial) was counterbalanced between subjects for both the choice RT tasks and the rhythmic tapping task.

#### Procedure

The experiment was delivered via online platforms, with participants using a web browser on their personal desktop or laptop computer to complete the required tasks. Participants accessed the study via Qualtrics, a secure online survey tool (qualtrics.com), where they first indicated they met the study criteria, digitally signed a consent form, and provided demographic information. Participants were then redirected to the Gorilla Experiment Builder (Anwyl-Irvine et al., 2020) (www.gorilla.sc) to enter the online experiment environment. Participants could not progress to Gorilla unless they were using a desktop or laptop computer with a keyboard.

In Gorilla, participants were given instructions on how to correctly perform trials of the 4-finger bimanual choice RT task. Participants performed the 2 versions of the task (symbolic stimuli/spatial stimuli) in consecutive blocks, with the order of task version counterbalanced between participants. There were 4 stimulus-response mappings for the symbolic and spatial versions of the task (**Figure 1A**). Before starting each block of trials, participants were given an opportunity to practice the task, to ensure they understood the instructions and to build familiarity with the stimulus-response mappings. Participants had to demonstrate that they could complete 6 consecutive practice trials without making an error before they were allowed to advance to the experiment. For each task version, participants performed 4 blocks of 15 task trials (15 per each 4 stimulus-response mappings) with an opportunity for a break every block. The order of the 4 choice RT stimuli was randomized between participants and between task versions. A random intertrial interval of 2-4 s was used so that participants could not anticipate stimulus onset timing (**Figure 1B**). Reaction time and the identity of keys pressed were recorded on each trial.

After completion of the bimanual RT task, participants began the 4-finger bimanual rhythmic tapping task. Participants were first shown instructions which described the types of movements that would be required. Instructions were accompanied by videos demonstrating the difference between symmetric and asymmetric tapping movements. After viewing the instructions, participants were subject to an attention check to ensure the instructions were properly read. They were asked how many taps they will perform per block. Participants that failed the attention check were looped back to the instruction screen and asked to read the instructions again. Participants were given no instructions on what type of movement strategy to use.

Upon passing the attention check, participants performed 4 versions of the 4-finger bimanual rhythmic tapping task (spatial-symmetric/spatial asymmetric/symbolic symmetric/symbolic asymmetric). Before each task version, participants practiced the task before beginning testing. During practice, cartoon hands were added to the task stimuli to explicitly demonstrate which fingers should respond to each stimulus. Participants were not allowed to progress to the first block of the task until they demonstrated they were capable of performing an abbreviated block of tapping (6 taps total) at a movement frequency of 1 hertz (Hz) for the given tapping condition. For each task version, participants performed 12 blocks of 32 taps (**Figure 1B**). Before the start of each block, a fixation cross was displayed with a clock graphic counting down from 3 seconds to allow participants to anticipate the start of the block. There was an opportunity for a break after every block. The movement frequency of rhythmic tapping started at 1 Hz and increased by 0.2 Hz each block, finishing at 3.2 Hz. For each stimulus type condition, participants always performed the symmetric version of the task first directly followed by the asymmetric version of the task. The order of the stimulus type condition was counterbalanced between participants and mirrored the order used in the 4-finger bimanual RT task. Key press timing and the identity of key presses were recorded in each task block.

#### Data processing

Custom MATLAB scripts were used to process data downloaded from Gorilla. For the 4-finger bimanual choice RT task, the RT for each responding finger was processed for each trial, and the mean of these 2 values was calculated to create a bimanual RT value for each trial. Trials were designated as an error if the wrong pair of keys were pressed, if no keys were pressed, if only one key was pressed, or if more than 2 keys were pressed. These trials were not included in RT calculations.

For the 4-finger bimanual rhythmic tapping task, an array of key press timing values was filtered from the raw data file for each finger and for each block. These arrays were used to calculate a relative phase value for each key press made. Relative phase values were calculated according to the formula:

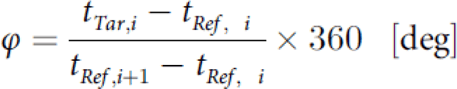

Here φ equals relative phase, t_tar,i_ equals the time of the *i*th tap of the target digit, t_ref,i_ equals the time of the time of the *i*th reference tap, and t_ref,i + 1_ equals the time of the *i*th +1 reference tap (Carson, 1995; Kodama et al., 2015) (see **Figure 2** for a graphical example). For symmetric blocks, the reference finger was the same effector on the opposite hand, while for asymmetric trials, the reference finger was the opposite finger of the opposite hand. The first 4 key presses were excluded from relative phase calculations, to allow participants time to get in rhythm.

**Figure 2.**
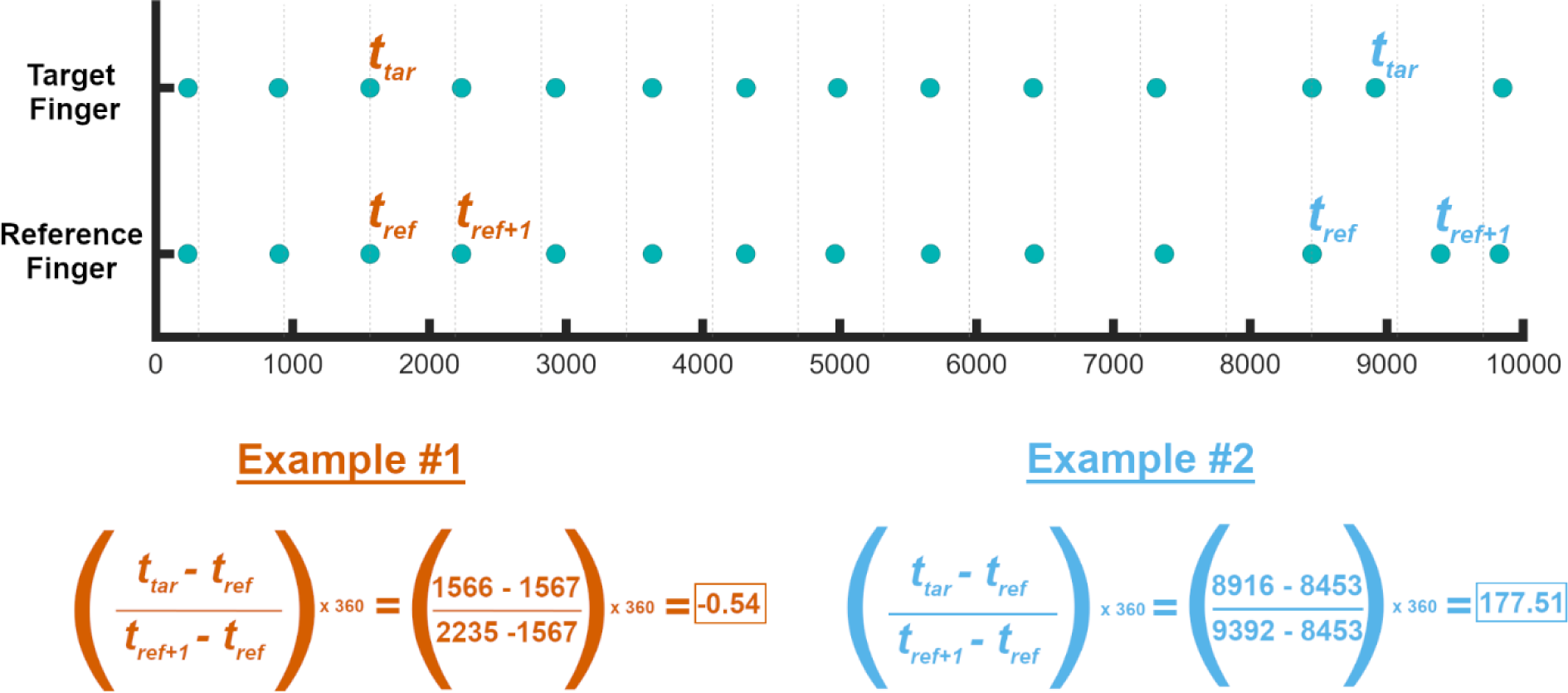
Method to calculate relative phase of bimanual coupling from key press data. An array of bimanual key press times from one task block is shown for target and reference fingers. The difference in time between each target finger key press and the temporally nearest reference key press is compared to the difference in time between that reference finger key press and the following reference finger key press. When bimanual temporal coupling is high, relative phase values are low, as is shown in example #1. When bimanual temporal coupling is low, relative phase values are high, as is shown in example #2.

#### Dependent variables

##### 4-finger bimanual choice reaction time task

- *Reaction time* – mean RT for symmetric and asymmetric trials was calculated for each stimulus type condition to provide a measure of RT magnitude. Error trials were not included in mean calculations.
- *Reaction time variability* – the standard deviation (SD) of RT for symmetric and asymmetric trials was calculated for each stimulus type condition. Error trials were not included in SD calculations.
- *Error rate* – the percentage of symmetric and asymmetric trials that were errors was calculated for each stimulus type condition.
- *Temporal coupling* – the mean absolute time between bimanual key presses for symmetric and asymmetric trials was calculated for each stimulus type condition.

##### 4-finger bimanual rhythmic tapping task

- *Tapping timing accuracy* – The mean absolute timing error was calculated from relative phase to provide a measure of tapping accuracy. Absolute timing error was calculated for each key press by subtracting the relative phase of each key press from ideal relative phase (0°) and calculating the absolute value of the resultant integer. The mean of absolute timing error was then calculated for each block.
- *Tapping stability* – The SD of absolute error was calculated for each block to provide a measure of tapping stability.
- *Transition occurrence –* Blocks in which a transition key press occurred (i.e., a key press with absolute relative phase greater than 180°) were marked as having a transition occurrence (1), whereas blocks in which no transition key press occurred were marked as having no transition occurrence (0).

#### Statistical analysis

Statistical analyses were carried out in RStudio. Post hoc analyses were performed using Bonferroni’s correction for multiple comparisons where appropriate.

##### Quality control

Since the experimental tasks was delivered online and there was no experimenter present to supervise performance, it is possible that participants did not properly understand the tasks or were not engaging in the task in the desired manner (e.g., making minimal effort, not paying attention, or performing the tasks incorrectly). To account for this possibility, quality control criteria were applied to the data before statistical analysis. For the 4-finger bimanual choice RT task, participants were removed from analysis if they produced errors at a rate of 3 SDs above the group mean error rate. This led to the removal of 10 participants from the dataset (n = 231). For the 4-finger bimanual rhythmic tapping task, participants were removed from analysis if their observed movement frequency was 20% above or below the desired movement frequency of 32 taps per block on at least 12 out of 48 task blocks, or if the grand mean of their mean absolute timing error scores was above 60°. This led to the removal of 49 participants from the dataset (n = 192).

##### 4-finger bimanual choice reaction time task

Before performing inferential statistics, mean RT, SD of RT, error rate, and temporal coupling values were assessed for skewness using the Shapiro-Wilks test. Results from these tests indicated that all variables were significantly positively skewed. To reduce the potential impact of skewness biasing results, linear mixed effect regression (LMER) analysis, rather than traditional *F*-tests, were used. LMER is robust against violations to distributional assumptions required by parametric tests (Schielzeth et al., 2020). LMER also accounts for random variation due to participants, through modeling of multiple intercepts for each participant as a random effect, rather than a single mean intercept (Baayen et al., 2008; Boisgontier & Cheval, 2016; Magezi, 2015; Nimon, 2012).

To test the effect of symbolic and spatial cueing stimuli on symmetric and asymmetric bimanual choice RT (Hypothesis 1), an LMER with fixed effects of STIMULUS TYPE (symbolic/spatial) and RESPONSE TYPE (symmetric/asymmetric) was used. Participant ID was included as a random effect. Separate LMERs were performed for mean RT, SD of RT, error rate, and mean temporal coupling.

##### 4-finger bimanual rhythmic tapping task

Before performing inferential statistics, mean absolute timing error and SD of absolute error values were assessed for skewness using the Shapiro-Wilks test. Results from these tests indicated that both variables were significantly positively skewed. To reduce the risk of data skewness biasing statistical models, observations that were deemed extreme outliers (i.e., 3 SDs above or below the mean) were removed from the dataset. The potential impact of skewness was also decreased by the decision to use LMER.

LMERs were performed to establish the moderating effects of within-subject fixed factors STIMULUS TYPE (symbolic/spatial) and RESPONSE PATTERN (symmetric/asymmetric) on the relationship between within-subject fixed continuous variable MOVEMENT FREQUENCY (1, 1.2, 1.4, 1.6, 1.8, 2, 2.2, 2.4, 2.6, 2.8, 3, 3.2 Hz) and mean absolute error and the SD of absolute error (Hypothesis 2, Hypothesis 3). Participant ID was included as a random effect.

Visual inspection of group means indicated the potential for a non-linear exponential relationship between MOVEMENT FREQUENCY and both mean absolute error and the SD of absolute error. Therefore, for these dependent variables, 2 LMERs were performed, one in which dependent variable data were untransformed and a second in which dependent variable data were log transformed to capture potential exponential components within the data. Models were compared using the Akaike information criterion (AIC) to assess whether a linear or exponential fit was more appropriate for each dependent variable.

Given that transition occurrence was a dichotomous variable (0 or 1), logistic regression was used to model the probability of transition occurrence across the continuous factor of MOVEMENT FREQUENCY and across categorical factors STIMULUS TYPE and RESPONSE PATTERN. Logistic regression is a statistical technique used to model dichotomous outcome variables(LaValley, 2008). This is achieved by modeling the log odds of the outcome as a linear combination of the predictor variables.

### Experiment 2

#### Participants

A total of 20 participants completed Experiment 2 (35.2 ± 11.3 years old; 14 female, 6 male; 16 right-handed, 4 left-handed). Twelve of these participants had previously participated in Experiment 1. Since the primary goal of Experiment 2 was to assess changes in movement quality generally across all response patterns and stimulus type conditions, a sample size of 20 was adjudged to highly powered enough to detect the predicted effect, given that each participant repeated the behavioural task of interest 4 times. The same inclusion criteria were applied as those used in Experiment 1. All protocols were approved by Behavioural Research Ethics’ Board at the University of British Columbia. Before participating, individuals provided written informed consent in accordance with the Declaration of Helsinki.

#### Experimental Design

The design of Experiment 2 was identical to Experiment 1, except participants only performed the 4-finger bimanual rhythmic tapping task in a laboratory setting. In addition, video of the participants hands was captured during task performance.

#### Procedure

The 4-finger bimanual rhythmic tapping task was performed as described in Experiment 1 on a desktop computer (iMac; Apple, CA) with a wired keyboard. Before participants began the task, a GoPro Hero 9 video camera (GoPro, CA) was placed below the computer monitor facing the participant to record finger movements from a frontal perspective. The camera lens was set to “narrow mode”, video resolution was set to 1080p, and video frame rate was set to 120 frames per second.

#### Data processing

Task data downloaded from Gorilla was processed as described in experiment 1. An open source deep learning toolbox (DeepLabCut (Mathis et al., 2018)) was used to estimate the 2D coordinate of index and middle fingers on every frame of the collected videos. DeepLabCut facilitates the training of a deep neural network to identify features of interest within every frame of a video. Labels were placed at the midpoint on the edge of the fingernails of the index and middle fingers of each video frame selected for neural network training. Labelled video frame data from every participant were included in the training dataset. Each participant contributed 60 video frames: 20 from a slow pace block, 20 from a medium pace block, and 20 from a fast pace block. Training frames were selected using a k-mean clustering approach that automatically identified representative frames. A single human rater labelled each training frame. ResNet-50 – a pretrained Convolutional Neural Network for image classification – was used as the initial weights of the neural network. The deep neural network underwent 600,000 iterations of training. Every frame of collected video was then processed through the trained network, producing 2D coordinate data for each finger on every frame of video, as well as a probability value indicating the networks confidence in the coordinate data for each frame. 2D coordinate data was smoothed across time with a gaussian kernel with a width of 3 frames.

Each participant’s 2D coordinate data was cut into task blocks using timestamp data acquired from Gorilla. Kinematic analyses focused on motion in the Y-plane of the 2D video, given that the majority of motion occurred in this dimension. To examine the effect of task conditions on relative kinematics, Y-plane coordinate data from each block was normalized to the time of the 1 Hz task blocks, i.e., 32 s or 3840 frames. This was achieved by interpolating Y-coordinate data for each block into the length of the first task block using the “interp2” function in MATLAB. A linear function was used to join interpolated data points. After interpolation, the “findpeaks” function in MATLAB was used to locate the relative timepoints in each block when peaks in the Y-coordinates occurred for each finger (i.e., the relative timepoints when key presses occurred). Using these indices, an average shape of finger trajectories adjacent to key presses was calculated for each finger and for each block, by calculating the mean of Y-coordinate data in a 120 frame window around the peak indices (see **Figure 6A** for an example). Peaks in which there were at least 12 frames where the DeepLabCut probability tracking rating were below 0.95 were not included in mean calculations, to reduce the effect of inconsistent tracking on trajectory shape.

#### Dependent variables

The dependent variables used to assess performance on the 4-finger bimanual rhythmic tapping task were the same as those described in Experiment 1. In addition, the following dependent variables were derived from mean peak shapes to test for changes in relative kinematics across task conditions:

- *Mean relative speed*: the relative speed of each finger in a time window of 120 frames around each peak was calculated by differentiating Y-coordinates with respect to time using the “diff” function in MATLAB. The mean of the resulting array of speed values was calculated for each tracked finger. The mean of these four finger values was calculated to give a single mean relative speed value for each task block.
- *Mean relative acceleration*: the relative acceleration of each finger in a time window of 120 frames around each peak was calculated by differentiating peak speed values with respect to time using the “diff” function in MATLAB. The mean of the resulting array of acceleration values was calculated for each tracked finger. The mean of these 4 values was calculated to give a single mean relative acceleration value for each task block.

#### Statistical analysis

Statistical analyses were carried out in RStudio. Statistical approaches to analyzing finger bimanual rhythmic tapping task performance data were the same as those described for experiment 1, except that LMERs were run only on log transformed mean absolute timing error and SD of absolute timing error values.

LMERs were performed to establish the effects of within-subject fixed factors STIMULUS TYPE (symbolic/spatial) and RESPONSE PATTERN (symmetric/asymmetric) and within-subject fixed continuous variable MOVEMENT FREQUENCY (1, 1.2, 1.4, 1.6, 1.8, 2, 2.2, 2.4, 2.6, 2.8, 3, 3.2 Hz) on relative range of motion, relative speed, and relative acceleration (Hypothesis 4). Participant ID was included as a random effect.

Visual inspection of group means indicated the potential for a non-linear exponential relationship between MOVEMENT FREQUENCY and relative acceleration. Therefore, for relative acceleration two LMERs were performed, one in which dependent variable data were untransformed and a second in which dependent variable data were log transformed to capture potential exponential components within the data. Models were compared using the AIC to assess whether a linear or exponential fit was more appropriate for each dependent variable.

## Results

### Experiment 1

#### 4-finger bimanual choice reaction time task

*Mean reaction time* – significant main effects of STIMULUS TYPE (ß = 192, 95% confidence interval [CI; 178 206], *p* <0.001, *Cohen’s d* = 0.87) and RESPONSE TYPE (ß = 50, 95% CI [39 61], *p* <0.001, *d* = 2.0) were found, as well as a significant STIMULUS TYPE x RESPONSE TYPE interaction (ß = 54, 95% CI [41 67], *p* <0.001; **Figure 3**). Estimated marginal means were calculated to further interrogate the interaction effect. A significant difference in mean RT was found for trials cued with symbolic stimuli and spatial stimuli, but the difference in mean RT was greater when trials were symbolically cued (µ = 104 ± 6, *p* < 0.0001, *d* = 1; symmetric: 747 ± 137 ms, asymmetric: 850 ± 178 ms) compared to those that were spatially cued (µ = 50 ± 6, *p* < 0.0001, *d* = 0.77; symmetric: 554 ± 101 ms, asymmetric: 605 ± 131 ms).

**Figure 3.**
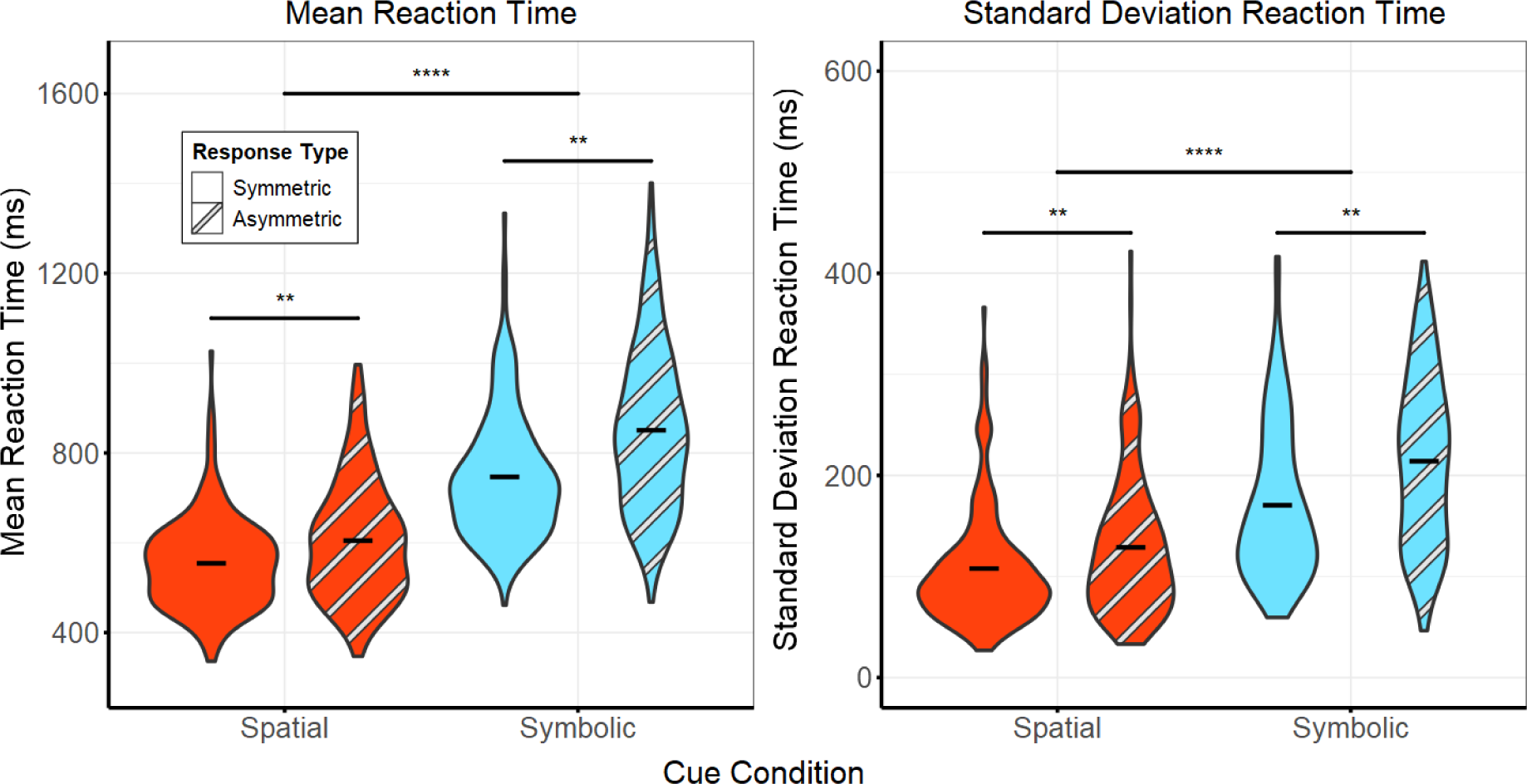
Bimanual 4-choice reaction time task results. Mean (left) and standard deviation (right) of reaction time (RT) values for symmetric and asymmetric response types collected in the spatially cued and symbolically cued bimanual 4-choice reaction time tasks. RT was significantly faster when movements were cued with spatially congruent cues and when movements were asymmetric. Furthermore, the difference in symmetric and asymmetric movements for the magnitude and variability of RT was significantly greater when movement was cued with abstract symbolic cues.

*Standard deviation of reaction time* – significant main effects of STIMULUS TYPE (ß = 63, 95% CI [54 72], *p* < 0.001, *d* = 1.0) and RESPONSE TYPE (ß = 20, 95% CI [13 29], *p* < 0.001, *d* = 0.5) were again shown, as well as a significant STIMULUS TYPE x RESPONSE TYPE interaction (ß = 23, 95% CI [11 34], *p* < 0.001, **Figure 3**). Estimated marginal means were calculated to further interrogate the interaction effect. A significant difference in SD of mean RT was found for trials cued with symbolic stimuli and spatial stimuli, but the difference in mean RT was greater when trials were symbolically cued (µ = 44 ± 4, *p* < 0.0001, *d =* 0.62; symmetric: 170 ± 75 ms, asymmetric: 214 ± 83 ms) compared to those that were spatially cued (µ = 21 ± 4, *p* < 0.0001, *d* = 0.37; symmetric: 108 ± 57 ms, asymmetric: 129 ± 67 ms).

*Error rate* – Higher error rates for symbolic cued movements and asymmetric movements were confirmed by significant main effects for STIMULUS TYPE (ß = 1, 95% CI [0 1], *p* < 0.001, *d* = 0.21; spatial: 2%, symbolic: 2.9%) and RESPONSE TYPE (ß = 1, 95% CI [1 2], *p* < 0.001, *d =* 0.33; symmetric: 1.7%, asymmetric: 3%). There was no significant STIMULUS TYPE x RESPONSE TYPE interaction effect for error rate.

*Temporal coupling* – Asymmetric responses were associated with a decrease in temporal coupling, as evidenced by significant main effect of RESPONSE TYPE (ß = 2, 95% CI [1 3], *p* < 0.001, *d* = 0.31; symmetric: 11 ms, asymmetric: 14 ms). There was no significant effect of STIMULUS TYPE on temporal coupling.

#### 4-finger bimanual rhythmic tapping task

##### Mean absolute timing error

Because the LMER fit to log transformed data produced a model with lower AIC (6368) compared to the LMER fit to untransformed data (72534), suggesting an exponential fit, we report results based on these transformed data. Mean absolute error increased exponentially with increasing movement frequency for all task conditions, as was evidenced by a main effect of MOVEMENT FREQUENCY (ß = 0.74, 95% CI [0.71 0.78], *p* <0.001), as shown in **Figure 4**. The detrimental effects of symbolic cues compared to spatial cues on mean absolute error increased exponentially with movement frequency, as was evidenced by a significant MOVEMENT FREQUENCY x STIMULUS TYPE interaction (ß = 0.19, 95% CI [0.15 0.24], *p* <0.001). Similarly, the detrimental effects of asymmetric patterns compared to symmetric patterns on mean absolute error increased exponentially with movement frequency, as was evidenced by a significant MOVEMENT FREQUENCY x RESPONSE PATTERN interaction (ß = 0.12, 95% CI [0.08 0.17], *p* <0.001).

**Figure 4.**
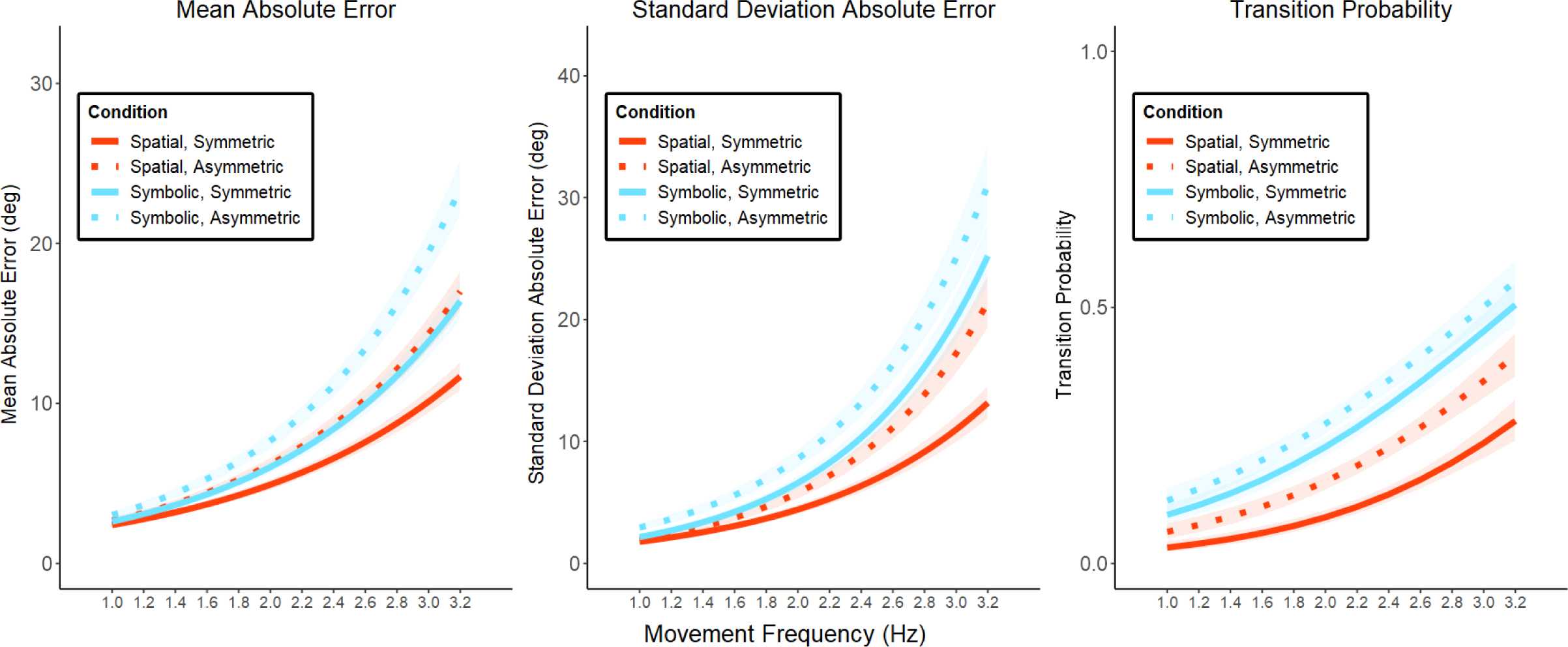
Bimanual 4-finger rhythmic tapping task results. Predicted task performance values in the rhythmic bimanual tapping task derived from linear mixed effect and logistic regressions. Lines represent predicted values and shaded regions represent 95% confidence intervals. An exponential increase in the mean absolute error of relative phase (left) and standard deviation of absolute error of relative phase (middle) was found. Effects of cue type (spatial vs. symbolic) and response type (symmetric vs. asymmetric) were low at slower movement frequencies and high at faster movement frequencies. The probability of transition out of the required response pattern also increased with movement frequency (right), but the effects of response type and cue type were consistent across movement frequencies.

##### Standard deviation of absolute timing error

Based on the LMER fit to log transformed data; the SD of absolute timing error increased exponentially with increasing movement frequency for all task conditions, as was evidenced by a main effect of MOVEMENT FREQUENCY (ß = 0.96, 95% CI [0.91 1.01], *p* <0.001). The detrimental effects of symbolic cues compared to spatial cues on the SD of absolute error increased exponentially with movement frequency, as was evidenced by a significant MOVEMENT FREQUENCY x STIMULUS TYPE interaction (ß = 0.22, 95% CI [0.15 0.29], *p* <0.001). Similarly, the detrimental effects of asymmetric patterns compared to symmetric patterns on the SD of absolute error increased with movement frequency, as was evidenced by a significant MOVEMENT FREQUENCY x RESPONSE PATTERN interaction (ß = 0.16, 95% CI [0.09 0.23], *p* < 0.001). The relative detrimental effects of asymmetric responses patterns compared to symmetric response patterns was greater for spatially cued movements compared to symbolically cued movements, demonstrated by a STIMULUS TYPE x RESPONSE PATTERN interaction (ß = 0.4, 95% CI [0.18 0.62], *p* <0.001). All 2-way interaction effects should be considered in light of a 3-way MOVEMENT FREQUENCY x STIMULUS TYPE x RESPONSE PATTERN interaction, which indicated that the relative detrimental effects of asymmetric patterns compared to symmetric patterns on the SD of absolute timing error with increasing frequency was greater for spatially cued movements than for symbolically cued movements (ß = −0.25, 95% CI [−0.35 −0.15], *p* <0.001) (**Figure 4**).

##### Transition occurrence

Logistic regression analysis revealed that the probability of a transition occurrence increased with increases in MOVEMENT FREQUENCY, as demonstrated by a significant effect of MOVEMENT TYPE (ß = 1.12, 95% CI [0.95 1.28], *p* <0.001). Symbolically cued movements were associated with a higher probability of transition occurrences than spatially cued movements, as demonstrated by a main effect of STIMULUS TYPE (ß = 0.77, 95% CI [0.25 1.29], *p* <0.005). Similarly, symmetric patterns were associated with a lower probability of transition occurrences than asymmetric movement patterns, as demonstrated by a main effect of RESPONSE PATTERN (ß = −0.94, 95% CI [−1.61 −0.27], *p* <0.01) (**Figure 4**).

### Experiment 2

Analysis of key press data in Experiment 2 largely replicated those found in Experiment 1. There were some exceptions, likely driven by decreased statistical power.

#### Mean absolute timing error

As shown in Figure 5, results from Experiment 1 were replicated. A main effect of MOVEMENT FREQUENCY (ß = 0.74, 95% CI [0.71 0.78], *p* <0.001), as well as significant MOVEMENT FREQUENCY x STIMULUS TYPE (ß = 0.19, 95% CI [0.15 0.24], *p* <0.001) and MOVEMENT FREQUENCY x RESPONSE PATTERN (ß = 0.12, 95% CI [0.08 0.17], *p* <0.001) interactions were found. However, the 3-way MOVEMENT FREQUENCY x STIMULUS TYPE x RESPONSE PATTERN interaction effect was not significant (ß = −0.14, 95% CI [−0.31 0.03], *p* = 0.12).

**Figure 5.**
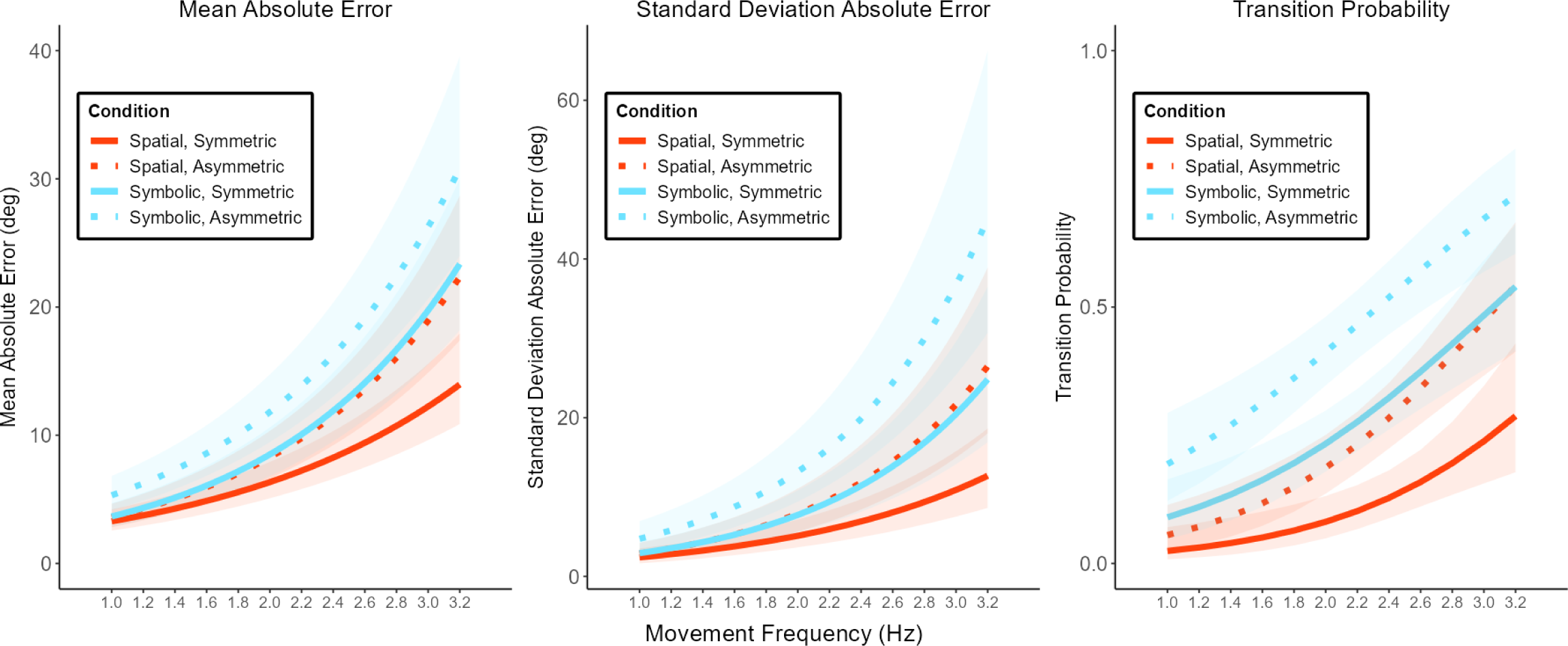
Experiment 2: bimanual 4-finger rhythmic tapping task results. Results from experiment 1 were largely replicated in experiment 2 with some exceptions. Notably, there was no main effect of response type on the probability of transition, although there was a non-significant trend (*p* = 0.11).

#### SD of absolute timing error

Results from experiment 1 were replicated. A significant main effect of MOVEMENT FREQUENCY (ß = 0.76, 95% CI [0.58 0.93], *p* <0.001) as well as significant MOVEMENT FREQUENCY x RESPONSE PATTERN interaction (ß = 0.25, 95% CI [0.03 0.48], *p* <0.001) were found. However, the MOVEMENT FREQUENCY x STIMULUS TYPE interaction was not significant (ß = 0.22, 95% CI [−0.01 0.44], *p* =0.059). Furthermore, the 3-way MOVEMENT FREQUENCY x STIMULUS TYPE x RESPONSE PATTERN interaction was not significant (ß = −0.2, 95% CI [−0.51 0.12], *p* = 0.22).

#### Transition occurrence

Results from experiment 1 were replicated with one notable exception. Significant main effects of MOVEMENT FREQUENCY (ß = 1.37, 95% CI [0.87 1.91], *p* <0.001) and STIMULUS TYPE (ß = 1.71, 95% CI [0.15 3.33], *p* <0.05) were found. However, the main effect of RESPONSE PATTERN was not significant, although it did approach significance (ß = - 0.75, 95% CI [−3.04 1.37], *p* = 0.11).

#### Relative speed

As shown in **Figure 6B** the relative speed of fingers increased with increasing movement frequency, (ß =0.18, 95% CI [0.15 0.22], *p*<0.001.

**Figure 6.**
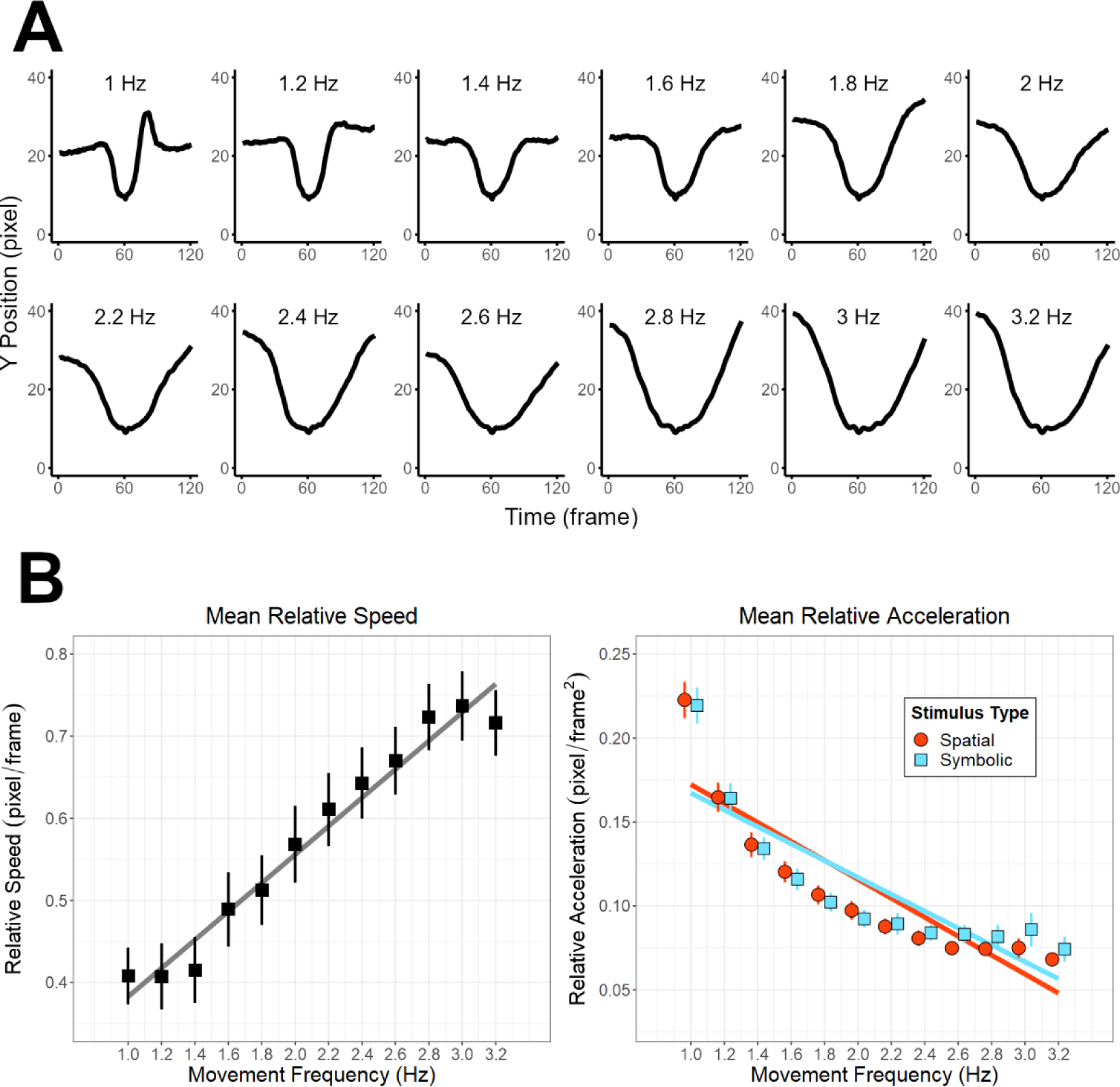
Increasing movement frequency was associated with a shift in movement quality. **(A)** Example of the average relative trajectory of the left middle finger in time windows around key presses in a single participant across movement frequency conditions. Trajectories were normalized by time to enable comparison of relative kinematics. **(B)** Mean relative speed (left) and mean relative acceleration (right) of all finger trajectories across movement frequency conditions. The mean relative speed of finger trajectories linearly increased with increasing movement frequency, while the mean relative acceleration linearly decreased with increasing movement frequency. Movements cued with spatially congruent stimuli showed a greater decrease in relative acceleration with increasing movement frequency. This indicates that the quality of finger kinematics transitioned from repeated discrete movements to smooth continuous movement as movement frequency increased.

#### Relative acceleration

The LMER fit to untransformed data produced a model with lower AIC (−3881) compared to the LMER fit to log transformed data (98), suggesting a linear fit. Therefore, we report results from the untransformed model. Relative acceleration of fingers decreased with increasing movement frequency as was evidenced by a main effect of MOVEMENT FREQUENCY (ß =-0.6, 95% CI [−0.6 −0.5], *p*<0.001). Relative acceleration decreased at a slightly lower rate with increasing movement frequencies for symbolically cued blocks compared to spatially cued blocks, as was evidenced by a significant MOVEMENT FREQUENCY x STIMULUS TYPE interaction (ß =0.1, 95% CI [0 0.1], p<0.05) (**Figure 6B**).

## Discussion

The current study demonstrated that the perceptual quality of visual stimuli used to cue movement timing has powerful effects on the control of discrete and rhythmic bimanual finger movements. In a bimanual 4-choice RT paradigm, actions cued with spatially congruent stimuli resulted in faster and less variable RTs compared to actions cued with abstract symbolic stimuli. Furthermore, the difference in RT between asymmetric responses versus symmetric responses was significantly reduced in the spatially congruent condition (**Figure 3**). These results largely mirror what has been found in bimanual reaching paradigms (Diedrichsen et al., 2001, 2003, 2006), providing evidence that crosstalk within response selection processes is the dominant source of interference constraining the planning of discrete asymmetric bimanual movements. As predicted, rhythmic bimanual movements became less accurate and less stable with increasing movement frequency; this relationship was best modeled as a non-linear exponential curve (**Figure 4**). Asymmetric movement patterns were less accurate and less stable than symmetric patterns. Similarly, when rhythmic movement frequency was cued with spatially congruent stimuli, accuracy and stability increased compared to conditions in which movement frequency was cued with abstract symbolic stimuli. The magnitude of these effects was minimal at lower frequencies but large at higher frequencies. Surprisingly, asymmetry effects were consistent across cueing conditions, suggesting that spatially congruent stimuli do not have the capacity to reduce asymmetry costs to temporal coupling of rhythmic bimanual movements. Increases in movement frequency were associated with increased mean relative speed and a decreased mean relative acceleration of finger trajectories, suggesting a shift in movement quality (**Figure 5**). A shift in movement quality might explain the non-linear fit between movement frequency and rhythmic task performance as well as the increase in magnitude of effect of response pattern type and movement cueing type across movement frequency.

### Spatially congruent cues reduce planning costs for asymmetric bimanual finger movements

As predicted, in a bimanual 4-choice RT task, responses cued with spatially congruent stimuli were faster and less variable than those cued by abstract symbolic stimuli, while symmetric responses were faster and less variable that asymmetric responses. Furthermore, the difference in RT from symmetric to asymmetric responses was greater for movements cued with abstract symbolic stimuli (**Figure 3**). These results extend findings from bimanual reaching paradigms and broadly supports the central processing hypothesis, which proposes that constraints in the production of discrete asymmetric bimanual movements stem from interference within central response selection processes that translate task relevant stimuli into action (Diedrichsen et al., 2001, 2003, 2006; Hazeltine et al., 2003). However, our results were not in complete agreement with bimanual reaching paradigms as there were significant differences between symmetric and asymmetric RTs in the spatially congruent cue condition. This discrepancy is most likely due to the increased statistical power in the current study, which had a much larger number of participants than previous bimanual reaching experiments and therefore better positioned to detect a small effect. The discrepancy with bimanual reaching experiments could also be due to differences in movement time between finger press and reaching movements. In reaching, movement time is on the scale of 100s of milliseconds, which might allow participants to extend motor planning into movement execution and make corrections online. Such an outcome is not possible in the task used in this study, because participants had to keep their fingers resting over the relevant keys during preparation which kept movement times very short. Another possible driver of discrepancy with bimanual reaching experiments is that the perceptual quality of stimuli contributed to discrepancies between studies using finger press vs. reaching movements. The spatially congruent stimuli used in the current study were presented at a ∼90-degree angle on a computer monitor facing the participant. In previous bimanual reaching studies, LEDs within the reaching space directly defined the required bimanual movement. It is possible that presenting spatially congruent cues at a ∼90-degree angle may have necessitated a mental rotation process which introduced a source interference that slowed down asymmetric responses in the spatially congruent condition.

### Increasing movement frequency drives a change in rhythmic movement quality from a series of discrete movements to continuous movement

When assessing how various task manipulations affect rhythmic bimanual task performance, it is important to concurrently track how the movement quality changes with movement frequency, because changes in movement quality can alter coordination principles (R. Ivry et al., 2004). When the kinematics of rhythmic bimanual movements comprise a series of discrete movements, temporal control is achieved through “event timing” representations (Robertson et al., 1999; Zelaznik et al., 2000, 2002, 2005). In this mode, spatiotemporal coupling is achieved by repeating movements at a set time interval, likely computed by the cerebellum (R. B. Ivry et al., 2002; Spencer et al., 2003). In contrast, when kinematics comprise smooth continuous movement, temporal control is achieved via “emergent timing” representations, where spatiotemporal coupling emerges via the control of higher order kinematic parameter such as speed, likely computed by transcallosal cortical networks (R. B. Ivry & Hazeltine, 1999; Kennerley et al., 2002; Robertson et al., 1999; Zelaznik et al., 2000, 2002, 2005).

We predicted that increasing movement frequency would necessitate a qualitative shift in kinematics from repeated discrete movements to smooth continuous movement. To investigate this hypothesis, we captured full spatiotemporal kinematics of task-relevant fingers during bimanual rhythmic task performance by applying deep learning algorithms to video recordings of finger movements. To quantify changes in kinematic quality, the mean relative speed and acceleration of finger trajectories during each task block was calculated. This analysis showed that the relative speed and relative acceleration of finger trajectories changed across movement frequency in a manner consistent with a shift from repeated discrete to continuous movement (**Figure 5**). Mean relative speed linearly increased with movement frequency, as the relative amount of time fingers spent resting on keys decreased, while mean relative acceleration linearly decreased with increasing movement frequency, as finger trajectories became smoother and more continuous with less dramatic shifts in relative speed. This change in movement quality provides important context with which to interpret temporal coupling results.

### Accuracy and stability of bimanual rhythmic finger tapping decreased exponentially with increasing movement frequency

As predicted, in *Experiment 1* the accuracy and stability of bimanual tapping and the probability of transition out of the desired movement pattern increased with movement frequency across all task conditions. The relationship between movement frequency and task performance variables were best modelled as exponential curves, with increases in movement frequency having a far greater effect on task performance variables at higher frequencies compared to lower frequencies (**Figure 4**). Given the finding that the quality of movement changed with increasing movement frequency, it seems likely that the exponential component within the data may have been driven by a change in coordination mode. At lower frequencies, participants likely favoured a strategy of rapid discrete bimanual movements separated by pauses, with pause length defined according to an event timing representation. Since results from the bimanual 4-choice RT task demonstrated that temporal coupling of discrete bimanual movements was high (∼11 ms on average), it follows that accuracy and stability of temporal coupling of bimanual rhythmic movements would be high when using a repeated discrete movement strategy. The use of a repeated discrete movement strategy could also explain how increases in movement frequency between low frequencies had a minimal effect on task performance. All that needed to be updated between blocks when using a repeated discrete movement strategy was the magnitude of the interval within the event timing representation, which is easily achieved by healthy participants when the intervals are regular (Semjen & Ivry, 2001; Yamanishi et al., 1980).

The exponential decrease in accuracy and stability shown in predicted values after approximately 2 Hz could be explained by a shift in strategy by participants towards smooth continuous movement. Within this coordination mode, instead of controlling movements according to an event timing representation, participants likely had to rely on a dynamic representation of speed derived from the flickering stimulus that denoted movement frequency. Given the relatively large decreases in task accuracy after this point, it appears that such a method of control was less effective for accurately coupling bimanual movements, and less capable of managing increases in target speed; increases in movement frequency was associated with large detriments to task performance.

### Effects of response symmetry and spatial congruence on rhythmic bimanual tapping stability emerge at higher movement frequencies

Changing the required response patterns from symmetric to asymmetric had the hypothesized effect on task performance; with symmetric patterns reducing the slope defining the relationship between movement frequency and tapping accuracy and stability, compared to asymmetric patterns. Whether the frequency of rhythmic movements was cued with repeating spatially congruent stimuli or abstract symbolic stimuli had a near identical effect on task performance.

Asymmetric response patterns and abstract symbolic cues were consistently associated with a greater probability of transition out of the desired response pattern across all movement frequencies. In contrast, the magnitude of the effects of response pattern type and timing cue stimulus type on bimanual coupling accuracy and stability was much greater at higher frequencies compared to lower frequencies.

Unexpectedly, the impact of response pattern symmetry on task performance for rhythmic movements was similar across cue types. Based on the central processing hypothesis, we predicted that spatially congruent cues would reduce the impact of asymmetric relative to symmetric response requirements on spatiotemporal coupling during the performance of rhythmic bimanual finger movements (Wenderoth et al., 2009). However, asymmetric response patterns were associated with similar detriments to performance across spatially congruent and abstract symbolic cueing conditions. One exception to this was in measures of stability of bimanual tapping. Here, an interaction effect was found between response pattern type and cue type. However, the direction of this interaction effect was the opposite to what was predicted, with symmetric and asymmetric instability increasing at similar rate across movement frequency for abstract symbolically cued response patterns but not spatially congruent cued response patterns.

Given that increasing movement frequency likely induced a qualitative shift in movement kinematics from repeated discrete movements to smooth continuous movement, it seems important to consider the increasing magnitude of effect of response pattern type and cue type across movement frequency. Since participants may have adopted a strategy of rapid discrete bimanual movements separated by pauses at lower frequencies, the task may have effectively mirrored the bimanual 4-choice RT paradigm. This could explain why cue type and response type had a relatively low impact on task stability at slow frequencies; results from the 4-choice bimanual RT task indicate that discrete bimanual movements have a high degree of temporal coupling regardless of cue (no difference) or response type (∼3 ms).

As movement frequency was gradually increased, at some point, participants changed movement strategy from repeated discrete bimanual movements to smooth continuous bimanual movements. Once a continuous movement strategy was selected, the basis for temporal control likely switched to from an internal representation of the time interval between events to an internal representation of speed(R. Ivry et al., 2004). Therefore, the capacity to accurately perceive speed from the flickering visual stimuli likely became a key component of task success.

Psychophysical studies of speed and flicker perception have demonstrated that both the contrast ratio (Blakemore & Snowden, 1999; Champion & Warren, 2017; Sotiropoulos et al., 2014; Stone & Thompson, 1992; Thompson et al., 2006; Thompson & Stone, 1997) and spatial frequency (Bowker, 1982; Brooks et al., 2011; Campbell & Maffei, 1981; Smith & Edgar, 1990) of flickering visual stimuli can affect the accuracy of perceived speed estimates. While the quality of the visual stimuli used in such studies are difficult to directly compare with the stimuli used to cue movement timing in the current study, the spatially congruent and abstract symbolic stimuli used to cue rhythmic movements did differ in terms of their contrast ratio and spatial frequency. Given the increased stability found in spatially congruent conditions, we assume that the combination of perceptual qualities of the spatially congruent stimuli facilitated a more accurate representation of speed over the abstract symbolic stimuli. Other researchers have shown that increasing cognitive load decreases the capacity to accurately perceive the presence of a flickering visual stimulus (Carmel et al., 2007). Therefore, it is possible that the greater cognitive load associated with the repeated translation of abstract symbolic stimuli had a compounding effect on the accuracy of perceived speed estimations in the abstract symbolic conditions, contributing to the decreased stability. Such a scenario could explain why stability decreases at a similar rate regardless of asymmetry in the abstract symbolic condition but not in the spatially congruent condition. The benefits of symmetric patterns on stability may have been overwhelmed by the compounding effects of increased cognitive load on an already noisier representation of speed. The compounding effect of cognitive load could also explain the ability to reduce relative acceleration with increasing movement frequency in the spatially congruent cueing condition compared to the abstract symbolic condition. A more reliable representation of speed may have allowed participants to more effectively smoothen bimanual rhythmic movements in the spatially congruent condition.

The finding that response pattern symmetry had a greater effect on temporal coupling at higher movement frequencies implies that the degree of interference in processes responsible for implementing asymmetric finger movements were also much greater when the quality of movement was continuous as opposed to discrete. The idea the movement quality drives coordination principles is important because while symmetric rhythmic bimanual movements have long been established as more stable than asymmetric rhythmic bimanual movements (Donchin & de Oliveira, 2004; Kelso, 1995; Swinnen, 2002; Swinnen & Wenderoth, 2004), previous investigations linking brain activity to bimanual finger tapping behaviour have not always explicitly considered movement strategy in putative brain-behaviour relationships (Liuzzi et al., 2011; Meyer-Lindenberg et al., 2002; van den Berg et al., 2010). Detriments to temporal coupling of continuous bimanual finger movements caused by response asymmetry is probably due to interference within motor processes responsible for implementing continuous asymmetric bimanual spatial trajectories. This form of interference seems to be independent to the interference caused by varying the perceptual qualities of movement timing cue, since performance detriments due to response pattern symmetry were mostly constant across spatially congruent and abstract symbolic cue types.

The neural source of response asymmetry interference may arise from transcallosal connections between the primary motor cortices, which have been shown to cause crossed facilitation between homologous effector representations under some movement conditions (Carson et al., 2004). In the case of asymmetric movements, where homologous muscles are active on asymmetric schedules, this could cause interference within effector representations which drives instability and inaccuracy in bimanual coupling. Another candidate is within systems responsible for defining spatial codes for movement. When these spatial codes are not perceptually symmetrical between hands, it may lead to interference between codes that reduces accuracy and stability of bimanual coupling. Studies with callosectomy patients indicate that such interference is probably localized within transcallosal circuits connecting the parietal cortices, since the capacity to couple bimanual reach directions is abolished only after callosectomy to the posterior third of the corpus callosum (Eliassen et al., 2000). Our data is not capable of distinguishing between these candidate neural sources of interference, since the asymmetric tapping conditions were both perceptually asymmetric and physically asymmetric in terms of muscle activation patterns. However, in a previous bimanual finger tapping task, perceptually symmetric movements were consistently more accurate than perceptually asymmetric movements, even when physically they were asymmetric (Mechsner et al., 2001). The finding that perceptual symmetry and not physical symmetry was the driver of the effect suggests that interference within systems representing movement at the level of spatial codes is the dominant source of interference contributing to detriments in tapping performance.

### Limitations

One limitation of the current study is that experiment 1 was conducted via online platforms, with participants performing the behavioural tasks at home on their own computers. Delivering the experiment via online platforms may have introduced uncontrolled environments with distracting background elements for some participants, which could have affected their performance. In addition, without the active supervision of an experimenter, participants may not have been motivated to perform the tasks to their best ability. These factors likely introduced noise into the dataset. However, the use of online platforms may have concurrently helped to mitigate these issues, since it facilitated the collection of a large sample size which reduced the impact of noise on results. The finding that results from Experiment 1 were largely replicated in Experiment 2, which took place in a controlled laboratory setting with a supervising experimenter, helps to further assuage fears on the validity of behavioural data collected via online platforms. Another limitation of the current study is that concurrent measures of unimanual task performance were not collected. This was driven by a desire to constrain the experiment time limit to 30 minutes to ensure participants gave full effort throughout the online session. The lack of a unimanual choice RT task or unimanual rhythmic tapping tasks limits our ability to assert whether the cognitive-perceptual constraints identified in this study generalize to unimanual movements or are specific to bimanual control. Future investigations could address this by additionally collecting unimanual versions of the described tasks. Finally, while we have discussed the likelihood that a change in the quality of movement (from discrete to continuous), alters the effect of stimulus type and response type on the stability of rhythmic bimanual finger movements, we cannot claim this relationship is causal, based on the data collected in this study alone. Future studies could better parse out causality by explicitly asking participants to adopt a repeated discrete movement strategy or a continuous movement strategy and examining the effect of these strategies on temporal coupling across a range of movement frequencies.

## Conclusion

Bimanual coordination is a complex phenomenon. Previous attempts to unify behavioural findings under a singular set of principles or map bimanual control to a specific set of brain regions have not fully appreciated the full diversity of the human bimanual repertoire and how small changes in the demands of tasks used to measure bimanual coordination can radically alter the underlying coordination principles at play. Therefore, calls to conceive of bimanual coordination as a phenomenon governed by a “coalition of constraints” (Carson, 2004; Carson & Kelso, 2004; Swinnen et al., 2004), with task demands modulating the relative contributions of given constraints and how they interact, seems to be a reasonable perspective. Such a view underscores that coordination principles will not always generalize between different movement types. This is highlighted in the current set of experiments, where we showed that increasing movement frequency can change the movement strategy selected by participants, which in turn appeared to transform the effect of response pattern asymmetry, as well as the effects of stimulus cue type on movement timing. Consideration of how changes in task demands prompt changes in movement strategies may be key to future efforts at uncovering principles of bimanual coordination and associated attempts at localizing the neural substrate which underpins their implementation.

## Acknowledgements

We extend thanks to Anna Zhu and Kaylee McGeough for their assistance in developing the DeepLabCut analysis pipeline used in Experiment 2.

## Data Availability Statement

The data that support the findings of this study are available on request.

